# An expansive internal chemosensory system in the insect abdomen implicates Gustatory Receptors in diverse aspects of interoception

**DOI:** 10.64898/2026.11.04.764383

**Authors:** Shinsuke Fujii, Hubert Amrein

## Abstract

Taste and olfactory receptors are gatekeepers that convey chemical information about the external environment. Their role in chemosensory systems has been extensively studied in several model system, which established their role in triggering discrete behavioral as well as physiological responses related to feeding and social interactions. Over time, cases of chemoreceptors expressed internally in organs not linked to the external chemosensory systems of taste and olfaction have been reported. Here, we present the existence of a complex internal chemosensory system in the abdomen of *Drosophila melanogaster.* We show that the adult fruit fly harbors a complex internal chemosensory system, composed of about 80 to 100 abdominal chemosensory neurons (ACSNs) expressing approximately 20 different *Gustatory receptor* (*Gr*) genes, including all sugar receptor and several members of the less conserved bitter receptor genes. The cells expressing specific *Gr* genes are stereotypically arranged along the entire circumference of the abdominal body wall, in proximity of muscle and fat layers. The ACSN fall into at least nine different subtypes based on co-expression analysis, projection trajectories into the Abdominal Ganglion (AG) and location of cell somata. Identification of different subpopulation of ASCNs expressing discrete combinations of *Gr* genes indicates that these receptor complexes have new, previously not considered roles in interoception, including monitoring of different nutrients, detection of cell metabolites and their byproducts and chemicals associated with pathogens.

## Introduction

The ability to sense and interpret chemical cues from the environment is a fundamental requirement for survival, guiding essential behaviors such as foraging, feeding, mate selection, and fear ^1^. The fruit fly *Drosophila melanogaster* is a powerful model system for dissecting the molecular and neural basis of chemosensation due to its genetic tractability, stereotyped behaviors, and a relatively simple nervous system ^2^. The sense of taste (gustation) is the primary sensory modality for evaluating food and other soluble chemicals, playing a pivotal role for identifying nutrients, such as sugars, amino acids, fats and salts while simultaneously detecting potentially harmful or toxic compounds ^3^. Thus, the decision-making process — to eat or not to eat — is orchestrated by activation of sensory neurons expressing chemical receptors tuned to these chemicals, ultimately controlling motor output in feeding behaviors ^4^, as well as regulating physiological adaptations to changing food availability.

The *Drosophila* gustatory system is dispersed across peripheral appendages including the proboscis, legs and wings and organized in stereotypically arranged sensory bristles (taste sensilla), each housing between two to four of gustatory receptor neurons (GRNs). Many but not all GRNs are tuned to one or related taste modalities, such as fatty acid and sugars, or bitter compounds and high concentrations of sodium chloride ^3^. The molecular basis for GRNs specificity is brought about by chemosensory receptor proteins, primarily the Gustatory Receptors (GR) and Ionotropic Receptors (IR) ^5,6^. Sugars, for example, are recognized by a small, conserved GR subfamily encoded by 8 genes, referred to as sugar *Gr* genes, which are largely co-expressed in a single GRN of all taste sensilla. Mutations in most sugar *Gr* genes reduces both physiological and behavioral responses to subsets of sugar, while the deletion of all eight genes results in sugar blind flies ^7^. Furthermore, restoring expressing of certain pairwise combination of sugar GRs in sugar blind flies, rescues sugar responses to some sugars, while expression of single sugar *Gr* genes does not ^7^, indicating that sugar receptor function as multimeric complexes composed of at least two different GRs ^8–13^. A larger set of GRs, which are partially co-expressed in another GRN of most taste sensilla, detect diverse non-nutritious chemicals which suppress feeding behavior and are perceived as bitter by humans ^3,14^. Similar to sugar GRs, bitter GRs also function as multimeric receptor complexes, generally composed of three or even four different subunits ^14^. Consistent with these genetic arguments, Cryo-EM studies suggest that GRs form tetrameric complexes ^15^, and two electrode patch clamp recordings of Xenopus oocytes expressing Gr43a indicate that these complexes function as ligand gated ion channels ^16^, similar to olfactory receptors (ORs), which are evolutionarily related to GRs ^17^.

A second group of about 20 receptors belongs to the Ionotropic Receptor (IR) family, with different members mediating the taste of fatty, carboxylic and some amino acids, respectively. IRs are at least partially co-expressed with GRs in both sweet and bitter GRNs, and they too function as tetra-heteromeric receptor complexes. In contrast to sugar and bitter receptors, IR based taste receptors share two core subunits, IR76b and IR25a. They are almost completely co-expressed across different types of taste neurons and combine with tunning IR subunits, which provide ligand specificity ^18,19^.

While functions of many *Gr* and *IR* genes and overall neuronal organization of respective chemosensory neurons have been characterized in remarkable detail, a growing body of evidence indicates that taste receptors have non-canonical roles outside the chemosensory system. Non-canonical expression of *Gr* genes was first reported for members of the *Gr28* gene subfamily, several of which are expressed in both central and non-chemosensory peripheral neurons, as well as non-neuronal cells in a range of tissues ^20^, but their specific roles has remained enigmatic outside the taste system. The best studied, non-canonical function for a *Gr* gene is that of *Gr43a*, which is expressed in about eight neurons in the posterior superior protocerebrum ^21^. There, Gr43a monitors hemolymph fructose and controls satiety dependent feeding behavior. While *Gr43a* is expressed in a few tarsal sugar GRNs, it plays no major role in sweet taste perception of flies. Additionally, Gr43a is expressed in the gustatory system as well as the brain of larvae, where it functions both as a classic sugar receptor as well as an internal nutrient sensor for fructose ^22^.

The insect abdomen accounts for more than 50% of a fly’s weight and houses virtually all major organs. It holds a majority of the digestive and respiratory tracts and the fat body (the insect equivalent of the mammalian liver), most hemocytes (immune cells), the Malpighian tubules (insect kidney), the dorsal vessel (heart chambers), the reproductive system and a major muscle layer lining the abdominal body wall. Due to the open circulatory system of insects, internal organs are exposed to hemolymph (insect blood), which is rich in nutrients, hormones and metabolites. The hemolymph is also a major reservoir where toxins and pathogens can accumulate, either acquired through ingestion, mating (in females) or through injury. Thus, it is not surprising that the fly’s abdomen is innervated by a complex array of about 1000 sensory neurons, including the enteric nervous system that controls gut function in larvae ^23^ and a diverse population of other, poorly characterized neurons, including multidendritic neurons that tile the abdominal body wall ^24^. Two papers have described the existence of internal chemosensory neurons in the abdomen, one describing expression of several members of the *Gr28* subfamily found in neurons located in the abdominal body wall ^20^, while the second reported expression of several *Gr* genes, including sweet and bitter taste *Gr* genes ^25^.

Here, we report the existence of a complex internal chemosensory system and present a detailed neuroanatomical and molecular *Gr* gene expression map in the insect abdomen. Using a combination of binary expression systems (*Gal4/UAS* ^26^ and *LexA/lexA_op_*^27^) with hexameric GFP and mCherry reporters ^28^, we uncover intricate anatomical features of a highly organized chemosensory neuron network and map cell somata, dendritic features and axonal projection of about 20 *Gr* genes encoding sweet, bitter and atypical taste receptors. Our data indicate that these abdominal chemosensory neurons (ACSNs) fall into at least nine distinct subtypes, based on co-expression, axonal projection patterns, dendritic arborization and location of cell bodies in the abdomen. Our work provides a comprehensive blueprint of an internal chemosensory network that integrates diverse chemical signals from the hemolymph to maintain physiological homeostasis. We hypothesize that interoception mediated through different subsets of ACSNs in response to changes in nutrients, metabolites and toxins and chemicals associated leads to the activation of neural circuits to adequately responds to these challenges physiologically and behaviorally.

## Results

Creating an internal chemosensory map in the abdomen is challenging. The only known physical landmarks of sensory neurons are the approximately 1000 tactile hair sensilla covering the abdominal cuticle, each consisting of three non-neuronal cells and a single mechanosensory neuron. In contrast to the external taste organs, where taste neurons are associated with stereotypically arrayed sensilla, there are no known physical landmarks associated with internal chemosensory neurons of the abdomen.

The central strategy for creating a physical map of ACSNs relied on available *Gr-GAL4* and *Gr-LexA* transgenes and gene knock-ins driving expression of sensitive reporters (*UAS-6xGFP/RFP* and *lexop-10xGFP*). We first mapped axon projections to the abdominal ganglion (AG) of ACSNs expressing a given *Gr* gene using a whole CNS preparation. This analysis provided an initial assessment for classifying ASCNs into subtypes. Specifically, similar AG staining pattern of different *Gr* genes suggests co-expression in ACSNs, while distinct AG staining patterns suggests that respective genes are expressed in different ACSNs subsets. We then used a abdominal pelt preparation in which ACSNs expressing a given *Gr* gene could be physically mapped onto an abdominal segment along the ventro-latero-dorsal axis. Finally, this analysis was complemented by co-staining experiments of flies carrying a *Gr-GAL4* and a *Gr-LexA* driver with respective reporters, resulting in a refined classification of a total of nine different ACSN subtypes. The abdomen preparation of adult flies expressing individual *Gr-GAL4* drivers also revealed distinct dendritic arborization patterns of different to ACSN subtypes, with proximity to different tissue layers (epidermis, abdominal muscle, fat) along the ventrodorsal as well as the anteroposterior axes.

Park and Kwon surveyed expression of all *Gr* drivers for expression in abdominal neurons by mapping axonal projections to the AG and surveying dendritic appearance of respective neurons ^25^. Several key questions remained unaddressed in this study, answers to which are critical to form hypotheses as to the roles of different receptors. First, it remained unresolved which *Gr* genes are co-expressed in specific ACSNs, and whether such co-expression follows the logic established for the gustatory system; Specifically, are receptors of the same modality co-expressed in subsets of ACSN? Second, the extent to which ACSNs are stereotypically organized was not investigated in that study, or where the somata of these ACSNs reside in the abdomen. Third, proximity ASCNs processes to potential tissues was not investigated, leaving open the most important question, namely from which organs these neurons might receive input. To address these questions, we analyzed *Gr* genes using improved imaging methods by takes advantage of highly sensitive reporters and by performing co-expression analyzes combining different binary expression systems, and by developing an abdominal pelt preparation that allows detailed mapping of ASCN cell-bodies and dendrites within the abdomen.

### Projections of ACSNs to the AG

#### Sugar *Gr* genes

Confocal images of antibody stained, whole CNS preparations of males expressing GAL4 and LexA as gene knock ins or transgenes for all sugar *Gr* genes were generated using z-stacks. Except for *Gr5a^LexA^* and *Gr43a^GAL4^* (Figure 1A and 1B), all other sugar *Gr* genes (*Gr61a-GAL4*, *Gr64a^GAL4^*, *Gr64b^LexA^*, *Gr64c^LexA^*, *Gr64e^LexA^* and *Gr64f^LexA^*; Figure 1C to 1H) presented with highly similar AG staining. Axons of these 6 *Gr* genes enter the AG in bilateral symmetrical fashion close to the lateral edge and curve inwards toward the midline where they terminate in a central stripe in the anterior 2/3 of the AG (Figure 1C’-H’). We consistently observed approximately 10 tracks entering the AG. A sagittal view of the VNC shows that axon projections from GRNs of the hindlegs and ACSNs are clearly segregated, with the former terminating in the ventral part of the third neuromere, while the latter curving laterally towards the center of the AG (Figure 1I).

**Figure 1.**
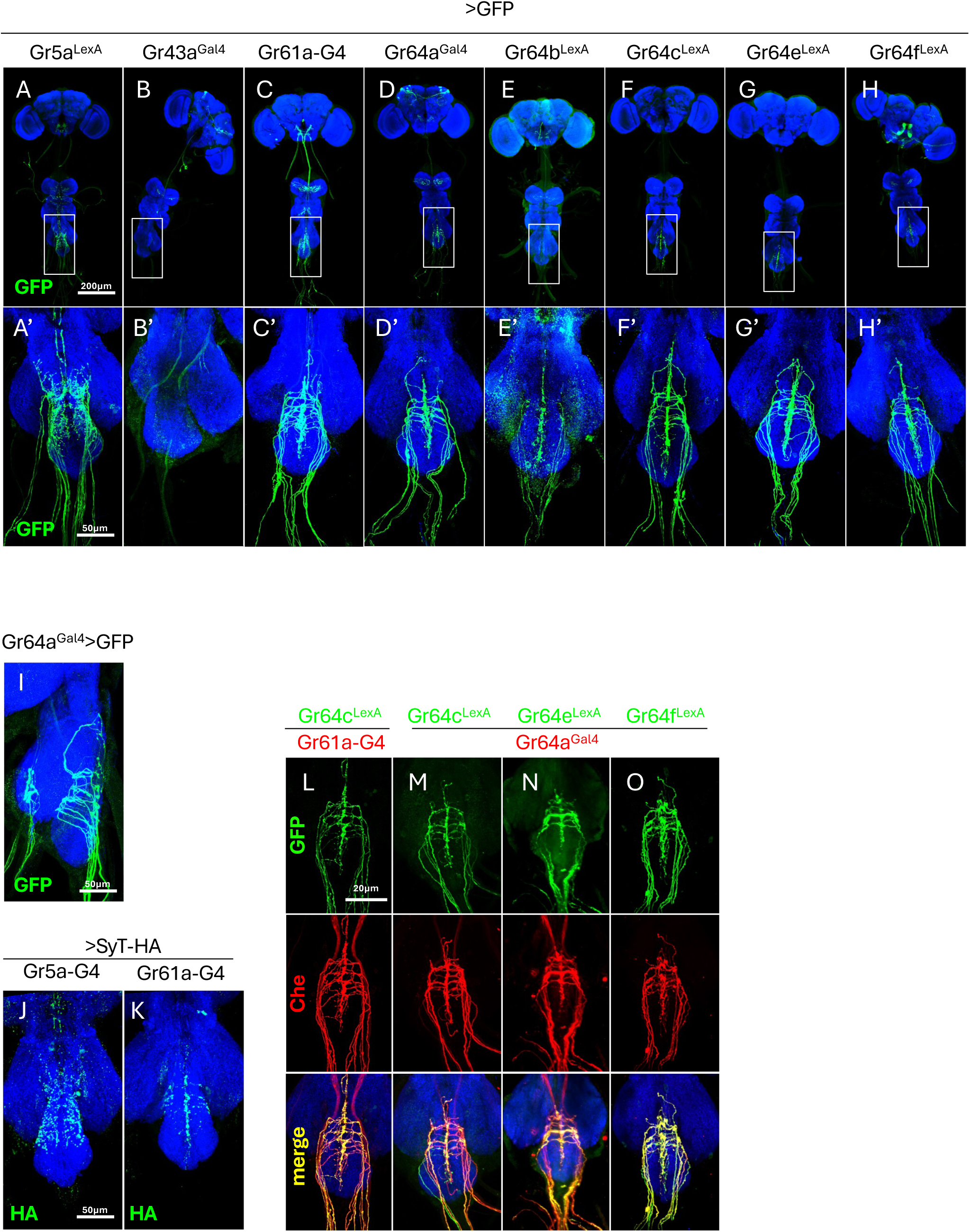
ACSNs of sugar Gr genes, except for Gr5a, exhibit highly similar projections to the AG. *Gr* gene expression in the male ventral nerve cord (VNC) was visualized with UAS- or LexAop-6xGFP. Panels A’–H’ are enlarged views of the AG from the panels A to H. (A) Neurons expressing Gr5a project to the lateral regions of the AG, avoiding the midline. (B) Gr43a expression is absent in males VNC. (C to H) The canonical sugar receptors (Gr61a, Gr64a, Gr64b, Gr64c, Gr64e, and Gr64f) show highly similar projection patterns, with axons converging on the dorsal midline of the AG. (I) A sagittal view of a *Gr64a^Gal4^* > UAS-6xGFP VNC, demonstrates that projections terminate in the dorsal-most layer of the AG. (J to K) Visualization of presynaptic terminals using a Synaptotagmin-HA reporter (UAS-syt-HA) confirms the distinct synaptic fields of Gr5a and Gr61a neurons. (L to O) Dual-labeling experiments confirm co-expression among canonical sugar Gr genes. Projections of Gr61a-Gal4 (driving mCherry) and *Gr64c^LexA^* (driving GFP) show extensive overlap (L). Similarly, projections from *Gr64a^Gal4^* (driving mCherry) show extensive overlap with those from (M) *Gr64c^LexA^*, (N) *Gr64e^LexA^*, and (O) *Gr64f^LexA^* (all driving GFP), indicating these genes are expressed in the same population of ACSNs. (P to Q) Projections labelled by Gr5a^LexA^ (gene knock in) and Gr5a-Gal4 (transgene) show broad overlap, suggesting that the transgene reproduces the gene-knock in expression.

*Gr5a^LexA^* neuron projections, while entering the AG on a similar trajectory as other sugar *Gr* drivers, are much more abundant, and in contrast to *Gr61-GAL4* neurons, their termini avoid the central midline (Figure 1A and 1A’). Segregation to different areas within the AG and hence, non-overlapping expression with the other six sugar *Gr* genes was supported by labeling presynaptic terminals using *UAS-SyT-HA* as a reporter: *Gr61a-GAL4* termini label mainly the central midline, while *Gr5a-GAL4* termini are excluded from that area and appear in a punctate array in the anterolateral area of the AG (Figure 1J and 1K). Furthermore, the total number of *Gr5a* axon tracks entering the AG is much larger (20-30) compared to other sugar *Gr* drivers. Finally, *Gr43a^GAL4^* showed no staining in the AG of males (Figure 1B), while in females, a unique projection pattern was observed (Supplementary Figure 1B), representing axon targets of neurons present in the uterus ^29^. No obvious sex-differences were observed for any of the other sugar *Gr* genes (Supplementary Figure 1A, 1C to 1H). In summary, the highly similar projection patterns of *Gr61a* and the *Gr64* expressing neurons suggests that they are co-expressed in ACSNs, while *Gr5a* appears to be expressed in different ACSNs. We observed complete overlap between the projection patterns of *Gr64c^LexA^*and *Gr61a-Gal4* in the AG (Fig. 1L). Similarly, *Gr64a^Gal4^* projections perfectly matched those of *Gr64c^LexA^*, *Gr64e^LexA^*, and *Gr64f^LexA^,* respectively (Fig. 1M-O).

#### Bitter *Gr* genes

We next characterize projections of ACSNs expressing bitter *Gr* genes in the abdomen (Figure 2). Many of these *Gr-GAL4* drivers produced, similar staining patterns, more restricted when compared to sugar *Gr* genes, with two pairs of axon tracts forming a heart-shaped trajectory along the lateral edge of the AG (Figure 2A, 2B, 2I, 2K, 2L, 2M and 2N). These axons coalesced along the AG midline. Altogether, seven bitter *Gr* drivers (*Gr8a*, *Gr22b*, *Gr32a*, *Gr39ab*, *Gr39b*, *Gr59a* and *Gr59d*) produced this highly similar projection patterns. In contrast, *Gr66a-GAL4* and *Gr89a-GAL4* labelled about 6 to 8 pairs of axon tracts that entered the AG more centrally, with elaborate punctate staining in the AG proper and labelling of the central midline only in the posterior part of the AG (Figure 2O and 2P). *Gr28b.d* and *Gr33a* showed a projection pattern similar to that of the 7 bitter *Gr* genes but appears extended to include at least one more pair of tracks in the anterior region of the AG (Figure 2G and 2J). Finally, three *Gr28* genes (*Gr28a*, *Gr28b.b* and *Gr28b.c*) show more extensive labeling that extends along the central midline of the VNC to the mesathoracic ganglion and includes some cell bodies within the AG (Figure 2C, 2E and 2F). We did not observe labeling for *Gr28b.a*, *Gr28b.e*, and *Gr94a-GAL4* (Figure 2D, 2H, 2Q). Gr94a-Gal4 shows weak and stochastic expression in AG.

**Figure 2.**
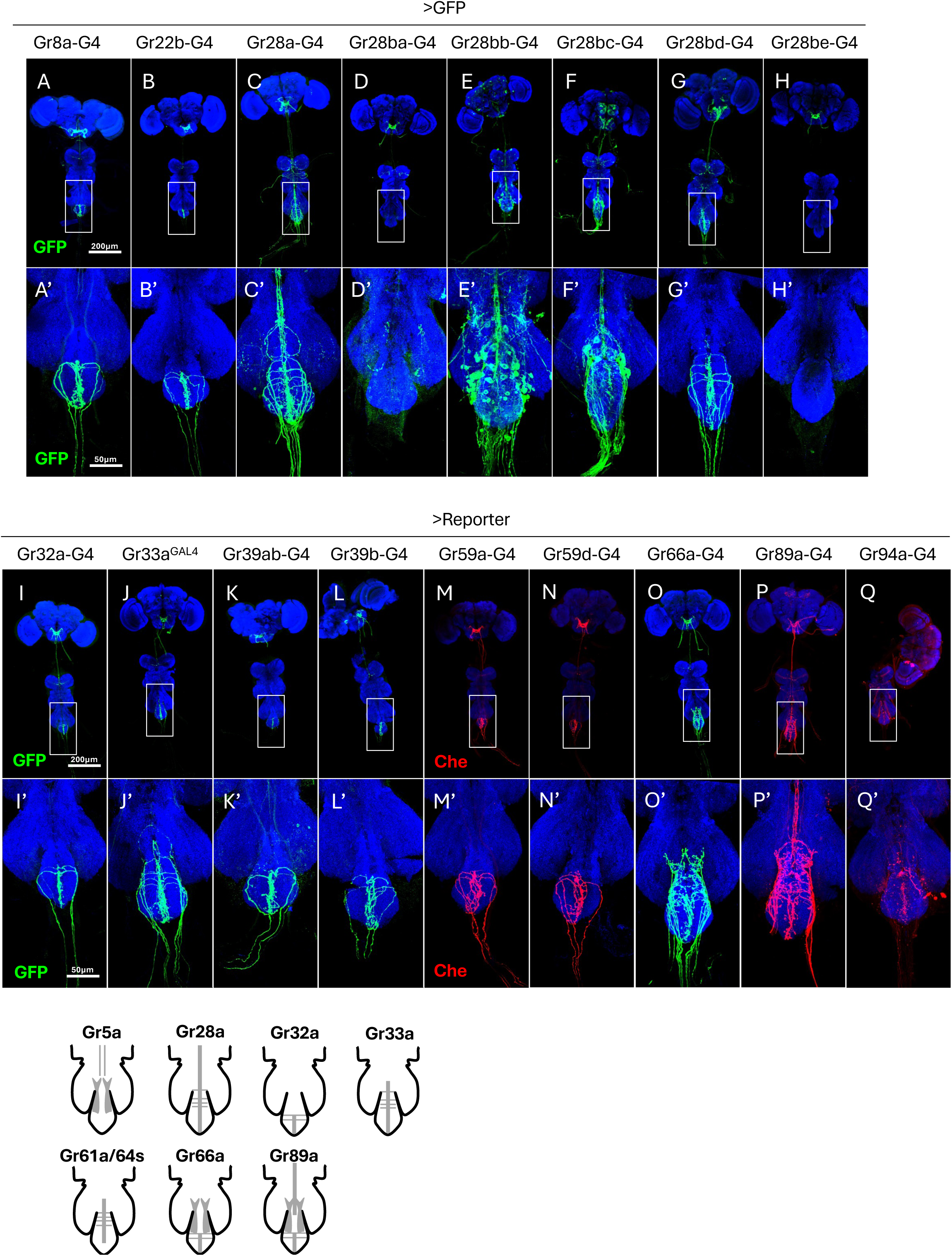
Projection patterns of bitter-responsive Gr genes in the AG. *Gr* gene expression in the male ventral nerve cord (VNC) was visualized with UAS- or LexAop-6xGFP. Panels A’–Q’ are enlarged views of the AG from the panels A to Q. (A, B, I, K, L, M, N) Several Grs including the pheromone/bitter receptor *Gr32a* labels a small number of axons that project to the midline and extend to the posterior tip of the ganglion. (C to H) The Ribonucleoside receptor *Gr28a* the related bitter *Gr28b* receptor genes of the Gr28 subfamily show distinct projection patterns (see text for detail). (J) The bitter co-receptor *Gr33a* labels neurons that also project along the midline. (O and P) canonical bitter receptor *Gr66a* and *Gr89a* labels projections that primarily target the lateral regions of the ganglion, a pattern similar to that of *Gr5a* (Figure 1A) in addition to midline and extend to the posterior tip. (Q) Gr94a shows no reproducible expression. (R) 7 typical patterns of the Gr projections in the AG.

We generated several pairwise combinations of *Gr* drivers using two bipartite systems simultaneously (GAL4 and LexA), which overall supported the largely mutually exclusive expression between *Gr5a* and the remaining sugar receptor genes, as well as the partial overlap of *Gr5a* and several bitter *Gr* genes (Supplementary Figure 2). However, since many of the ascending axons coalesce into bundles, the visualization of incoming axons cannot conclusively determine the extent of co-expression between *Gr* genes.

### Sensory neuron architecture in the abdomen

Most ACSNs are associated with trichoform sensilla, morphologically identifiable as sensory hairs located on the tergites. Since internal organs are largely devoid of sensory neurons, except for the reproductive organs and posterior gut/rectum, we suspected that ACSN somata and their dendritic networks are anchored within or near the abdominal body wall, as this is the sole tissue providing structural integrity and anatomical definition. Thus, we developed an abdomen preparation of the pelt and body wall (hairs, cuticle, body wall muscle and fat layer) that maintained the integrity of the neuroanatomical architecture (AG, including the soma, dendrites and axons of afferents; see Material and Methods). We first assessed the neural composition of the abdomen using the pan-neural marker *nSyb-GAL4*, driving expression of both cellular GFP and nuclear RedStinger which revealed a rich, largely stereotypical assembly of small cell bodies harboring the nucleus, bearing short visible processes (Figure 3A). These cells are bipolar mechanosensory neurons that send a dendrite to the base of a hair covering the abdomen (Figure 3B and 3C). In each segment, a prominent dorsoventral tract descends along the body wall and turns anterior towards the AG upon reaching the ventral midline region. A second axon bundle joins this prominent track near the lateral-ventral midline. Both minor and major tracts harbor four to six larger cell somata at defined positions, identifiable as yellow dots (co-localization of GFP and redStinger). In the ventral region, the segmental axon tracts converge into a thick fiber which contain two additional clusters of large cell somata/nuclei that ascends to the AG. In summary, neuroanatomical analysis of abdominal neurons revealed the existence of about 100 large sensory neurons that appear stereotypically arranged in a segment-specific manner, sending axons along the body wall toward AG.

**Figure 3.**
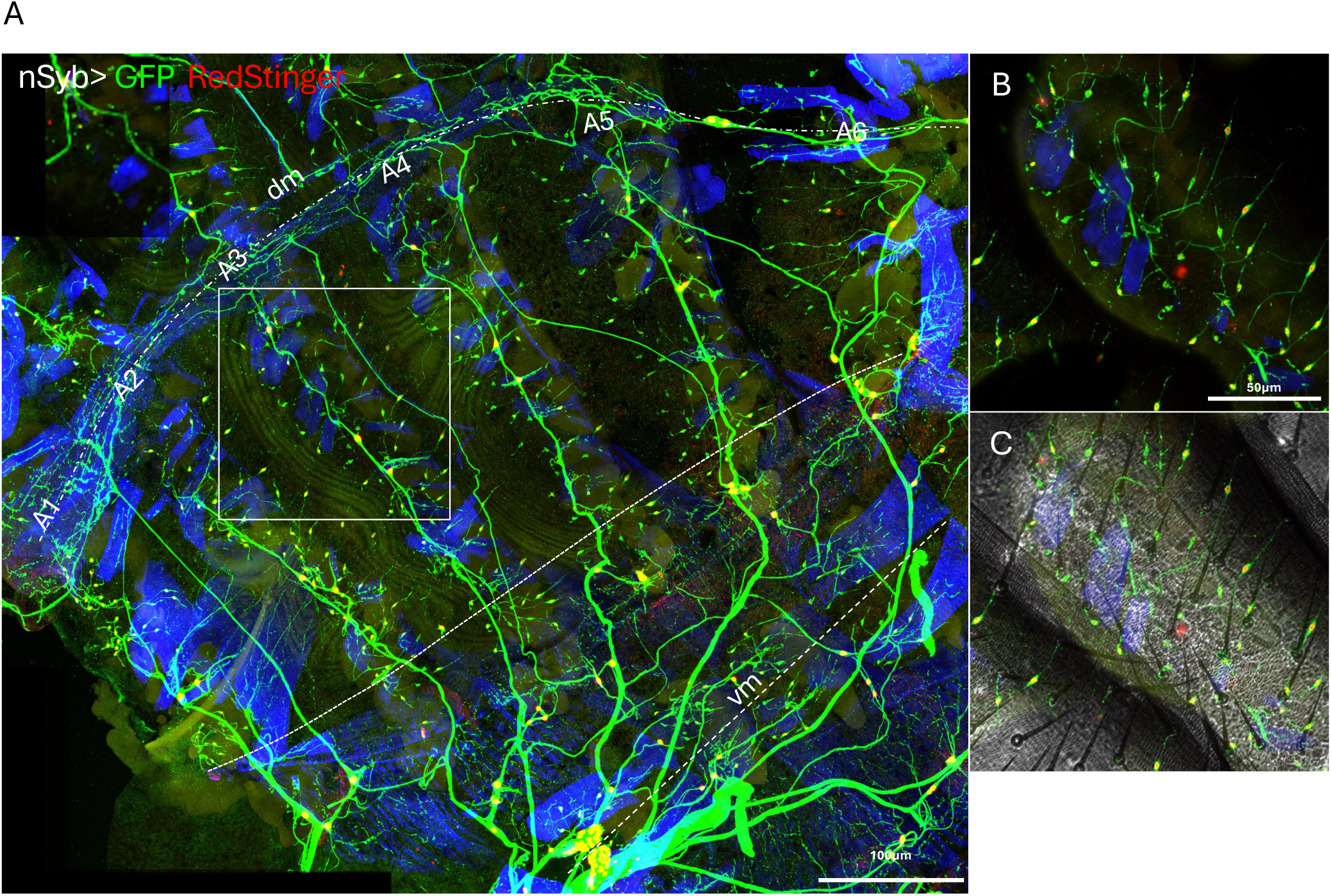
A Comprehensive Map of Internal Chemosensory Neurons on the Abdominal Body Wall. (A) Pan-neuronal labeling with *nSyb-Gal4*, driving expression of UAS-6xGFP (processes) and RedStinger (nuclei), reveals the entire network of neurons associated with the adult male abdominal body wall. (B-C) Spatial segregation of neuronal layers. Optical sectioning of the preparation shown in (A). The layer containing the fine-caliber dendrites of external sensory neurons (B). ACSNs are characterized by large somata and thick processes, which reside in a deep layer of the body wall (C).

As our analysis will show, more than half of these neurons are ACSNs expressing specific sets of *Gr* genes. We have been able to assign these ACSNs to one of nine subtypes, based on the location and set of *Gr* genes each of these ACSN subtypes expresses. Molecular signatures to ACSNs were obtained by performing antibody staining experiments of flies expressing single or pairs of *Gr* drivers and mapping respective neurons in the abdominal wall (Figures 5 and 6, Supplementary Figure 4), a summary of which is shown in Figure 4. We find that *Gr61a* and members of the *Gr64* cluster are all ventrally located, and co-staining of flies expressing *Gr61a* and *Gr64c*, as well as *Gr64a* and *Gr64c* indicates that these sugar *Gr* genes are co-expressed (Figure 5A and 5B; Table 1). In contrast, *Gr5a* neurons are found in three distinct regions across different regions in the abdomen: One subset of *Gr5a* neurons is located laterally, and a second is arranged in the dorsolateral region of abdomen, with each group having a single neuron/segment/side (Figure 6A to 6F). Similar to *Gr61a/Gr64*, there is also a *Gr5a* cluster in the ventral region, but co-staining of *Gr61a-GAL4* or *Gr64a^GAL4^* with *Gr5a^LexA^* shows that the latter is expressed in different group of neurons (Figure 5C and 5D, Table 1). This was further evident from analysis of dendritic arrays of *Gr5a* and *Gr61/Gr64* neurons (Supplementary Figure 3). The *Gr5a* dendrites are densely arrayed in the form of two lattice-like pattern in a band along the anterior posterior axis in the ventral and ventral-lateral region, respectively (Supplementary Figure 3A, 3B and 3E). The lateral *Gr5a* neurons generate a different dendritic network, reminiscent of type IV multidendritic neurons found in the larvae (Figure 6B) ^20^. *Gr61a/Gr64* dendrites appear less expansive and are exclusively restricted to the ventro-lateral region of the abdomen, intermingled with *Gr5a* lattice-type dendrites (Supplementary Figure 3C, 3D and 3F). Interestingly, these two neural processes reside in distinct layers of the fat body muscle layer, with the *Gr5a* neurons occupying the outer layer more densely and in contact with muscle tissue of the abdominal wall, and the *Gr61a/Gr64* dendrites the inner layer, closer to the fat cells (Supplementary Figure 3).

**Figure 4.**
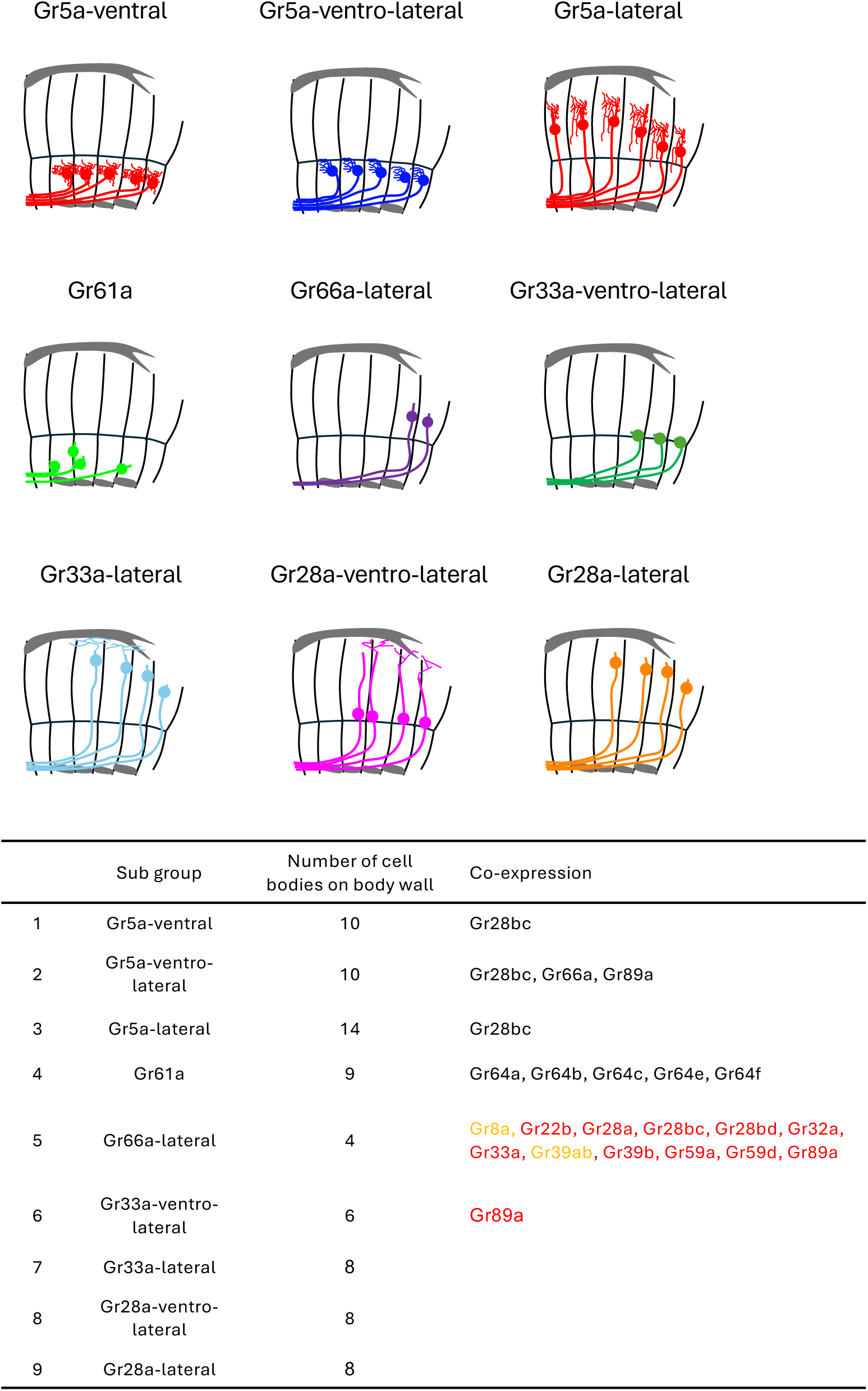
A schematic map summarizing the identity and location of the nine major ACSN subtypes subtypes. The map shows the right hemisphere of the abdominal wall. The gray object on top, aligned gray objects at the bottom in the map indicate dorsal tube and muscles at the ventral midline, respectively. Classification is based on receptor gene expression, projection pattern and soma position. Cell bodies (not to scale) are enlarged in size for enhancing visibility. Subgroup (1) Gr5a-ventral; (2) Gr5a-ventro-lateral; (3) Gr5a-lateral; (4) Gr61a; (5) Gr66a-lateral; (6) Gr33a-ventro-lateral; (7) Gr33a-lateral; (8) Gr28a-ventro-lateral; (9) Gr28a-lateral.

**Figure 5.**
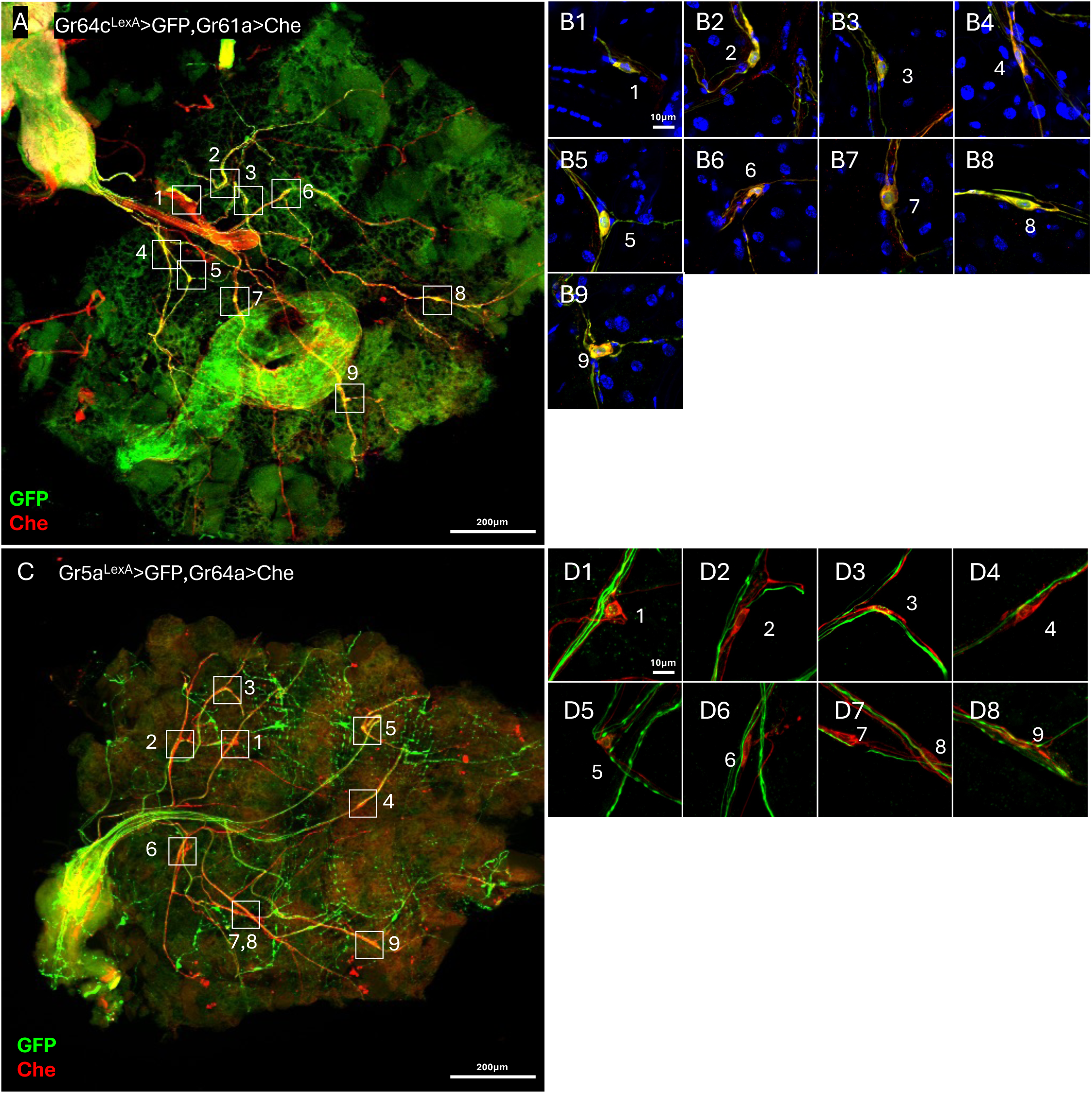
Peripheral co-expression analysis confirms sugar *Grs* and *Gr5a* mark distinct neuronal populations. Double-labeling of neurons on the ventral abdominal body wall was performed to confirm co-expression patterns observed centrally in the AG. All images are oriented with the anterior to the left and posterior to the right. (A) The canonical sugar receptors Gr61a and Gr64c are co-expressed in the same ACSNs. A male expressing GFP under control of *Gr64c^LexA^* (green) and mCherry under control of *Gr61a-Gal4* (red) reveals overlap in the cell somata (yellow). (B1 to B9) High-magnification views of individual somata, confirming co-localization of both reporters within the same cells (C) Gr5a and the sugar receptor Gr64a are expressed in distinct, non-overlapping neuronal populations. A male expressing GFP under control of *Gr5a^LexA^* (green) and mCherry under control of *Gr64a^Gal4^* (red) reveals that axon tracts of the two neuronal populations run in parallel but remain clearly segregated (D1 to D8) High-magnification views of individual somata confirm segregation at the cellular level. Somata are exclusively labeled with either GFP (Gr5a-positive) or mCherry (Gr64a-positive).

**Figure 6.**
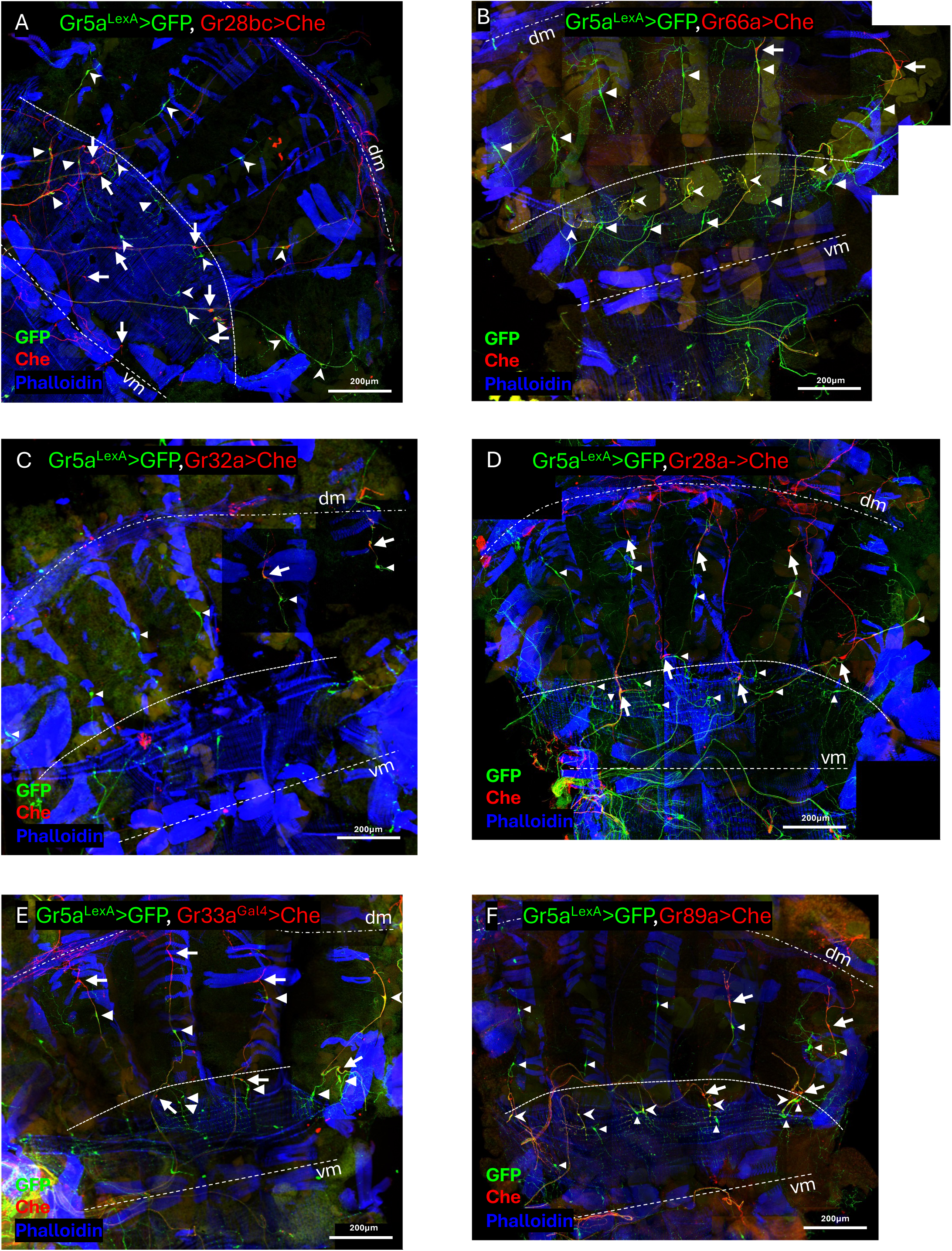
(A-F) The sugar *Gr5a* gene is co-expressed with bitter Gr genes in subsets of ACSNs. Neurons expressing *Gr5a^LexA^* only (green) are indicated by white triangles, while neurons expressing the indicated bitter Gr-Gal4 only (red) are indicated with a white arrow. Gr5a is co-expressed in some Gr28bc (A), Gr66a (B) and a few Gr33a (E) expressing neurons (yellow, white arrowhead). Many, but not all bitter Gr expressing and of Gr5a expressing neurons extend processes along the same track.

**Table 1.** Frequency of Grs co-expression in the abdominal neurons.

|  |  | *Gr5a<sup>LexA</sup>* | *Gr64b<sup>LexA</sup>* | *Gr64c<sup>LexA</sup>* | *Gr64f<sup>LexA</sup>* |
|---|---|---|---|---|---|
| Whole Ventral Body Wall | *Gr61a-G4* | 0.0/6.0 (n=2) | 5/8 (n=1) | 9.0/9.0 (n=3) | 9.5/9.5 (n=2) |
|  | *Gr64a<sup>Gal4</sup>* | 0/9 (n=1) |  | 9.0/9.0 (n=3) |  |
| Half Sagittal Body Wall and Whole Ventral Body Wall | *Gr22b-G4* | 0.0/2.0 (n=4) |  |  |  |
|  | *Gr28a-G4* | 0.0/8.0 (n=2) |  | 0/8 (n=1) |  |
|  | *Gr28bb-G4* | 0.0/2.0 (n=3) |  |  |  |
|  | *Gr28bc-G4* | 8.7/16.7 (n=3) |  |  |  |
|  | *Gr32a-G4* | 0.0/2.0 (n=4) |  | 0.0/1.7 (n=3) |  |
|  | *Gr33a<sup>Gal4</sup>* | 0.3/6.3 (n=3) |  | 0.0/6.5 (n=2) |  |
|  | *Gr39ab-G4* | 0.0/1.5 (n=2) |  |  |  |
|  | *Gr39b-G4* | 0.0/1.5 (n=4) |  |  |  |
|  | *Gr59a-G4* | 0/2 (n=1) |  |  |  |
|  | *Gr59d-G4* | 0.0/2.0 (n=3) |  |  |  |
|  | *Gr66a-G4* | 4.7/6.7 (n=3) |  |  |  |
|  | *Gr89a-G4* | 4.5/9.0 (n=2) |  | 0.0/8.5 (n=2) |  |
|  | *ppk-G4* | 17.0/17.0 (n=3) |  |  |  |

We next examined number and location of ACSNs expressing bitter *Gr-GAL4* drivers, and when appropriate, doing so along *Gr5a^LexA^* (Figure 6A to 6D; Supplementary Figure 4). Neurons expression bitter *Gr* drivers revealed several different subtypes. As expected from projection patterns to the AG, drivers for *Gr22b*, *Gr32a*, *Gr39b*, *Gr59a* and *Gr59d* labeled only a pair of laterally located neurons in segment 5 and 6 (Figure 6C; Supplementary Figure 4). Both these neurons are unique, in that they express by far the highest number of genes, including the three other bitter *Gr* genes (*Gr33a*, *Gr66a* and *Gr89a*; Figure 6E, 6B and 6F) as well as all three members of the *Gr28* subfamily (*Gr28a*, *Gr28bc*, and *Gr28bd* (Figure 6D and 6A, Supplementary Figure 4B). The second most diverse ACSNs are the five ventro-lateral neurons expressing *Gr5a,* which also express *Gr28bc*, *Gr66a* and *Gr89a-GAL4* (Figure 6A, 6E and 6F). Interestingly, *Gr28bc* is also found in the two other *Gr5a* expressing subtypes located in the ventral and lateral region of the abdomen. *Gr33a* defines two additional subtypes in the ventrolateral region (3 neurons, co-expressing *Gr89a*) and the lateral region (4 neurons), with no other examined receptor co-expressed in these ACSNs (Figure 6E). Lastly, we found that the known RNA/ribonucleoside receptor Gr28a is expressed in 3 ventrolateral and 4 lateral neurons (Figure 6D) in proximity of neurons expressing *Gr33a* (Figure 6E). Afferents of *Gr33a* and *Gr28a* neurons follow the same tracks towards the AG, and dendritic processes patterns are similar. However, it is currently not possible to unequivocally determine whether *Gr33a* and *Gr28a* are expressed in the same neurons or represent the same subtype or are different entities showing similar morphology.

We note three cases in which *Gr* expression diverges from the general feature described thus far: These includes *Gr43a*, exhibiting female-specific expression, with sensory neurons integrated in the reproductive tissue (uterus) ^21^, and at least two *Gr28b* genes (*Gr28b.b* and *Gr28b.c*), expressed in central neurons located in the AG. We previously reported that these *Gr28b* genes are broadly expressed across numerous central neurons in larvae ^30^.

## Discussion

The ability of an organism to sense its external environment is fundamental to survival, but of equal importance is the capacity to monitor its internal state—a process known as interoception. Our study provides a comprehensive anatomical and molecular blueprint of a sophisticated chemosensory system within the adult *Drosophila* abdomen, revealing a previously underappreciated network of internal sensory neurons. By systematically mapping the expression and projection patterns of key gustatory receptors, we demonstrate that the abdomen is a major hub for internal chemosensation, featuring distinct neuronal populations organized to monitor the fly’s internal chemical environment. Our findings challenge the view of gustatory receptors as purely external tastant receptors and establishes the fly abdomen as a powerful model for dissecting the principles of interoception.

A central discovery of our work is the functional divergence of two major neuronal systems in the abdomen: one that expresses the canonical carbohydrate receptor complex, and another marked by *Gr5a* that appears to be repurposed for a distinct, novel role. The first system comprises neurons that co-express the canonical ‘sugar GRs’ (*Gr61a, Gr64a-f*). These neurons project symmetrically to the dorsal midline of the AG, with cell somata located in the ventral region, and peripheral processes that appear to be able to sample the hemolymph as well as connecting with internal tissues, most notably the fat body — the fly’s equivalent of the mammalian liver and of importance for many metabolic processes. We hypothesize that these neurons function as homeostatic sensors, reporting the physiological status of fat cells and/or monitoring the hemolymph for fluctuations in key metabolites like trehalose and other circulating sugars. By conveying nutrient status information to the central nervous system, activation of this circuit could trigger compensatory adjustments in feeding behavior, nutrient storage, or metabolic processes, thus acting as a critical feedback loop for maintaining energy balance. This finding reframes the function of the well-known sugar GR complex, expanding its role from a simple detector of external food to a sophisticated monitor of internal energy reserves.

*Gr5a,* the remaining member of the sugar *Gr* gene subfamily, is expressed in an entirely different and significantly larger subset of ACSNs. In the gustatory system, this receptor is broadly co-expressed in sweet taste neurons along with other sugar receptors ^14^ and is required for the sweet taste of trehalose and other sugars ^31,32^. However, analysis of a *Gr5a-GAL4* transgene and especially *Gr5a^LexA^*, a *LexA* knock-in into the *Gr5a* locus, revealed that this receptor is expressed in additional non-sweet taste neurons in the main taste organs, the labial palps, suggesting that it has a function in the perception of other taste modalities. The finding that *Gr5a* is expressed in ASCNS not only devoid of other sugar *Gr* genes, but co-expressing at least one bitter *Gr* genes in multiple ACSN subtypes provides new evidence that this receptor has additional functions. We propose that GR5a functions as a key core subunit for diverse receptor complexes. While in the taste system, it co-operates with other sugar GRs subunits to form sugar receptors, in a ACSNs GR5a assembles with non-sugar GRs to form receptors for non-sugar compounds. Intriguingly, *Gr5a*-positive ACSN subtypes (1, 2 and 3) display a dendritic network reminiscent of multidendritic neurons present in the larvae, which is distinct form the dendritic signature of ACSNs expressing either the sugar *Gr* genes (ACSNs sub type 4; Figure 4) or ACSN subtypes characterized by expression of Gr66a, Gr33a and Gr28a (subtypes 5, 6, 7, 8 and 9; Figure 4). Notably, *Gr5a* positive in ASCNs co-express ppk (Supplementary Figure.4F and Table 1), a channel known to play a role in proprioception in larvae ^33^. This fact, and the observation that Gr5a dendritic processes appear to contact the abdominal body wall muscle (Supplementary Movie 1) suggests that GR5a and bitter GR co-expressed in these neurons have multisensory properties and play a role in both proprioception as well as interoception, although we cannot exclude the possibility that GR complexes in these neurons have acquired novel functions in proprioception.

Our investigations have broad implications for our understanding of interoception across the animal kingdom. Atypical expression of taste and olfactory receptors in internal tissues, such as the gut, airway, and brain, is a conserved phenomenon observed also in vertebrates, including humans ^34^. However, the functions of internally expressed chemosreceptors have been difficult to dissect in complex vertebrate systems, or systems with a limited repertoire of molecular-genetic tools. Our work establishes the *Drosophila* abdomen as a powerful and genetically accessible model to uncover the fundamental principles of how an organism monitors its internal chemistry. The anatomical blueprint we provide lays the essential groundwork for future functional investigations.

In conclusion, our work expands the concept GR proteins in *Drosophila* taste, moving it beyond the external gustatory organs. The presented study provides a foundational map that will guide future investigations into the mechanisms of interoception, transforming our understanding of how an animal senses not only the world around it, but also the world within.

## Materials and Methods

Fly strains used in this study were obtained from the following resources: UAS-6xGFP (BDSC_52262), UAS-6xmCherry (BDSC_52267, BDSC_52268), UAS-RedStinger (BDSC_90912), UAS-SyT-HA (BDSC_6924), lexAop-6xGFP (BDSC_52265), QUAS-6xmCherry (BDSC_52269), Gr5a-Gal4 (BDSC_57592), Gr8a-Gal4 (BDSC_57593), Gr22b-Gal4 (BDSC_57604), Gr32a-Gal4 (BDSC_57622), Gr33a (BDSC_31425), Gr39a.b-Gal4 (BDSC_57632), Gr39b-Gal4 (BDSC_57635), Gr59a-Gal4 (BDSC_57648), Gr59d-Gal4 (BDSC_57653), Gr61a-Gal4 (BDSC_57658), Gr66a-Gal4 (BDSC_57670), Gr89a-Gal4 (BDSC_57676), Gr94a-Gal4 (BDSC_57685), ppk-Gal4 (BDSC_32079), ppk-QF (BDSC_66475) (Bloomington Stock Center); Gr5a, Gr64a, Gr64b, Gr64c, Gr64e, Gr64f, Gr28a-Gal4, Gr28ba-Gal4, Gr28bb-Gal4, Gr28bc-Gal4, Gr28bd-Gal4 and Gr43a.

### Immunohistochemistry and Imaging

To visualize neuronal projections, fluorescent reporters (UAS-6xGFP, UAS-6xmCherry, lexAop-6xGFP, QUAS-mtdTomato) were expressed using appropriate Gal4, LexA, or QF driver lines. For nuclear labeling, UAS-RedStinger was used.

Central nervous systems (CNS) and abdomens from adult flies (7-14 days old) were fixed for 30 minutes in 4% paraformaldehyde in 1X phosphate-buffered saline (PBS) with 0.2 % Triton-X100 (PBST) at room temperature. Flies were washed three times in PBST and dissected in PBST. Tissues were blocked for 1 hour in PBST containing 5% heat inactivated goat serum at 4 C°.

Samples were incubated with primary antibodies overnight at 4°C. The following primary antibodies and dilutions were used: Chicken anti-GFP (1:10,000; Novus, NB100-1614), Rabbit anti-mCherry (1:10,000; Abcam, ab167453), Mouse anti-nc82 (1:50; Developmental Studies Hybridoma Bank) to label synaptic neuropil in the CNS. For synaptic terminal visualization (Fig. 1J, K), samples expressing *UAS-syt-HA* were stained with rabbit anti-HA (1:500; GeneTex, GTX115044).

After primary antibody incubation, tissues were washed four times in PBST and incubated with secondary antibodies overnight at 4°C. The following secondary antibodies and dilutions were used: Goat anti-chicken Alexa Fluor 488 (1:200; Thermo Fisher Scientific), Goat anti-rabbit Alexa Fluor 568 (1:200; Thermo Fisher Scientific), Goat anti-mouse Alexa Fluor 647 (1:200; Thermo Fisher Scientific), For abdominal body wall preparations, F-actin in muscle fibers was counterstained by Alexa Fluor 647 Phalloidin (1:500; AAT Bioquest, 23159) during the secondary antibody incubation. Following three times washes in PBST, samples were mounted on slides with VECTASHIELD® PLUS Antifade Mounting Medium with DAPI (Vector Laboratories).

Images were acquired using a Nikon A1R Confocal Microscope System. Unless otherwise specified (e.g., Fig. 1I), all CNS images were generated from Z-stacks scanned from the dorsal side. Image stacks were processed and assembled into maximum intensity projections using Fiji/ImageJ.

---

**Figure S1. Expression of sugar *Gr* genes in the female ventral nerve cord reveals a sexually dimorphic pattern for *Gr43a*.** To assess possible sexual differences, the expression patterns of the same set of sugar *Gr* genes (Figure 1) were examined in the adult female ventral nerve cord (VNC). Except for *Gr43a^GAL4^* (B), projection patterns of all sugar *Grs* are similar if not identical to that of males. *Gr43a*, which is absent in the male VNC (Figure 1B), shows robust and unique labeling in the female AG, confirming its previously reported sexual dimorphism. Representative images for each Gr line are shown.

**Figure S2. Co-expression analysis defines distinct chemosensory neuron projection patterns in the AG.**

(A to L) Double-labeling experiments were performed in the VNC to map the relationships between different chemosensory neuron populations. In all panels, enlarged images of the AG are shown (A’-L’).

(A-G) The sugar *Gr* population is distinct from other chemosensory neurons. The reference sugar Gr gene *Gr64c^LexA^* (driving GFP, green) was combined with Gr5a and indicated bitter Gr-Gal4 lines (driving mCherry, red).

As expected the Gr5a-Gal4 projections show largely overlap with *Gr5a^LexA^* projections.

(H to L) Neurons expressing Gr5a are a heterogeneous population that partially overlaps with some bitter-sensing neurons. The reference *Gr5a^LexA^* gene (driving GFP, green) was combined with indicated bitter Gr-Gal4 drivers (driving mCherry, red). Projections from members of the Gr28 subfamily and the sugar receptor Gr64a (G to K) are distinct from Gr5a projections, but a subset of Gr66a-Gal4 projections co-localizes with *Gr5a^LexA^* projections, appearing as yellow.

**Figure S3. Peripheral Arborization Patterns Reveal Distinct Terminal Targets for Gr5a and Sugar Gr Neurons.**

(A-F) Gr5a-positive neurons arborize extensively within the outer layers, whereas sugar Gr neurons do not. The ventral abdominal body wall of a male expressing GFP driven by *Gr5a^LexA^* (green) and mCherry by *Gr64^Gal4^* (red) is shown. At a superficial focal plane, Gr5a-positive neurons display a dense, lattice-like arborization pattern in the outer body wall, associated with muscle tissue (A). In contrast, Gr64a-positive neurons show minimal terminal branching in the same layer (C). At a deeper focal plane closer to the internal organs, the main axon bundles of both the Gr5a (B) and Gr64a (D) populations run in parallel pathways. 3D cross-sectional reconstructions (E, F) of the regions indicated by white squares in A-D. The reconstruction confirms that Gr5a arborizations (E, green) penetrate the outer layers corresponding to muscle, while Gr64a axons (F, red) are confined to the inner layer above the fat body.

(G, H1-H4) Canonical sugar Gr neurons terminate on distinct internal tissues, including the fat body. *Gr64c^LexA^*-positive neurons (driving GFP, green) were visualized alongside fat body cells marked by C7-Gal4 (driving mCherry, red). A wide-field view showing the distribution of Gr64c-positive axon terminals (green, white arrowheads; G). High-magnification views of a single terminal (H1-H4) from the region indicated by the white square in G. A maximum intensity projection shows the terminal ending (H1). Individual optical slices at different depths (H2-H4) reveal the intimate contact between the neuronal terminal (green) and the surface of a fat body cell (red).

**Figure S4. *Gr5a* neurons are distinct from most of other Gr expressing neuron populations.**

Neurons only expressing *Gr5a^LexA^* (green, white triangles) are distinct from neurons expressing Gr-Gal4 (red, white arrow). The cell bodies of the indicted bitter Gr genes are located in the lateral area of the last two segments but reside more dorsally compared to Gr5a neurons Four additional cell somata of Gr28bd expressing neurons (B) reside also more dorsally than more Gr5a cell bodies.

**Movie S1. Gr5a-positive neurons innervate the abdominal muscle layer.**

A three-dimensional visualization of the abdominal body wall from a Gr5a^LexA^ > LexAop-GFP fly. Gr5a-positive neurons are labeled with GFP (green), and muscle fibers are counterstained with phalloidin (red). The movie presents a Z-stack of confocal slices, scanning from the outer body wall inward. The fine terminal processes of the Gr5a neurons can be seen extending into and arborizing within the phalloidin-labeled muscle layer, demonstrating their intimate association with this tissue.

## Notes

### Competing Interest Statement

The authors have declared no competing interest.

### Funding

- NIH Common Fund: R01 DC018403-01A1

