## Supplemental Figures for "An expansive internal chemosensory system in the insect abdomen implicates Gustatory Receptors in diverse aspects of interoception"

### Supplementary Figure 1 (female CNS)

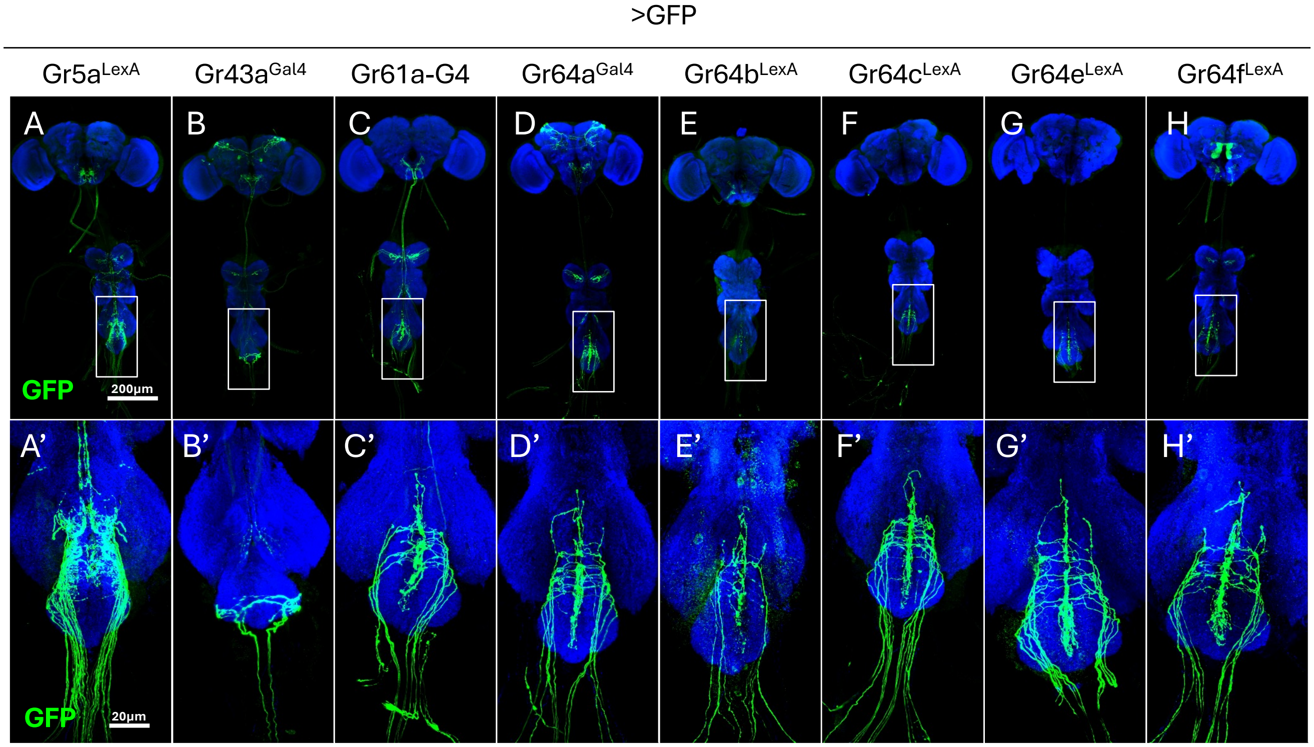

### Supplementary Figure 2

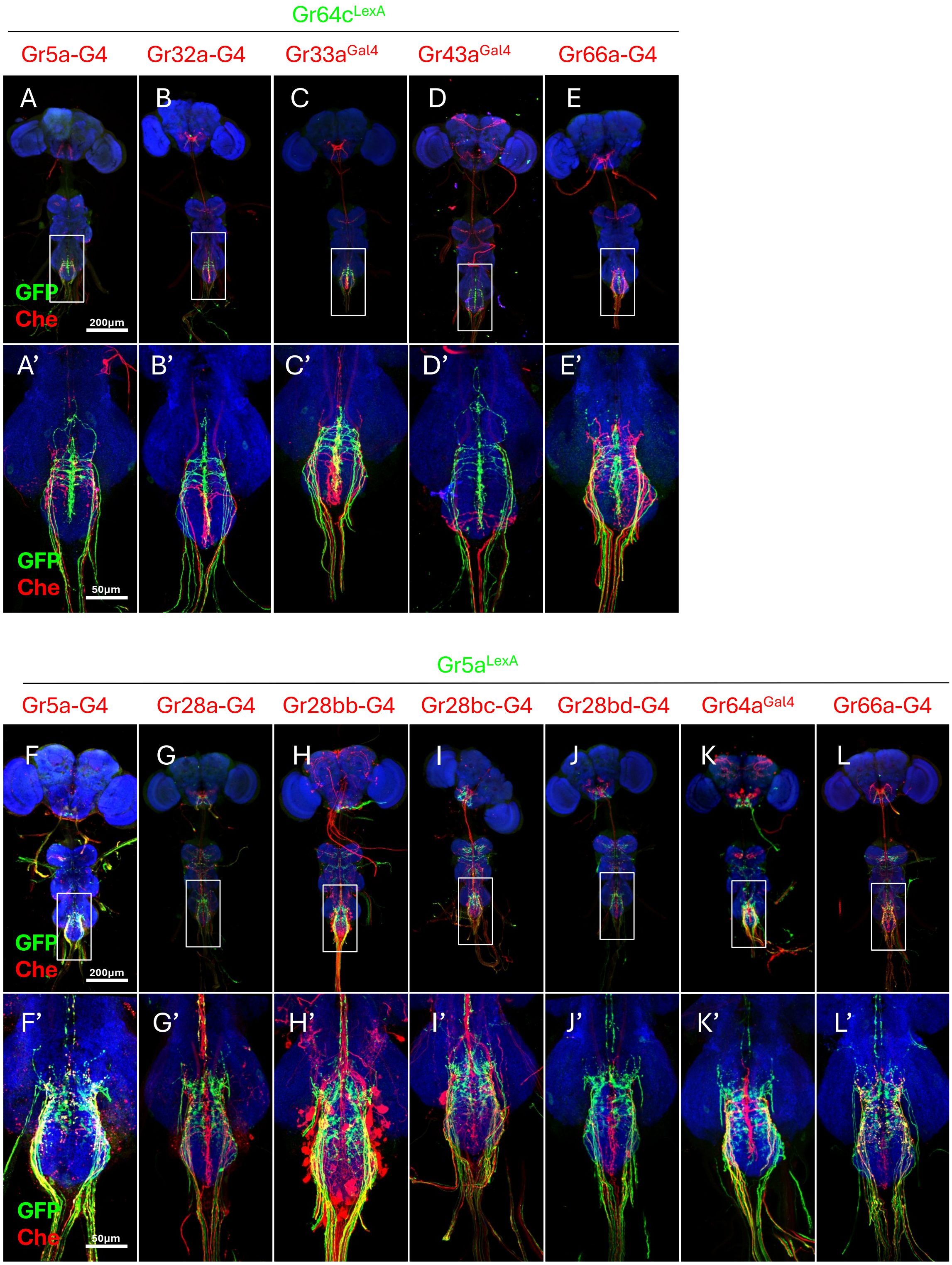

### Supplementary Figure 3

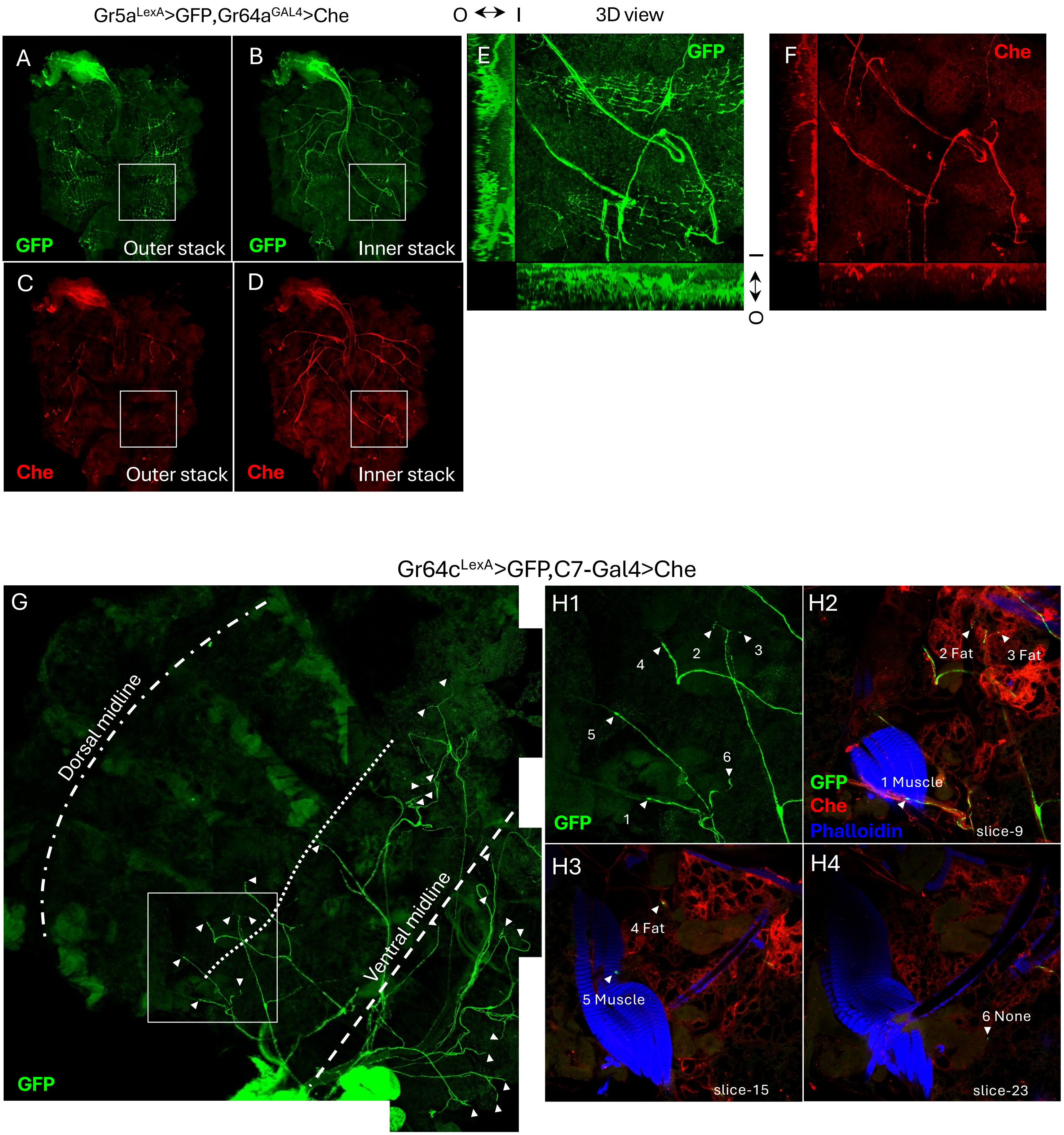

### Supplementary Figure 4

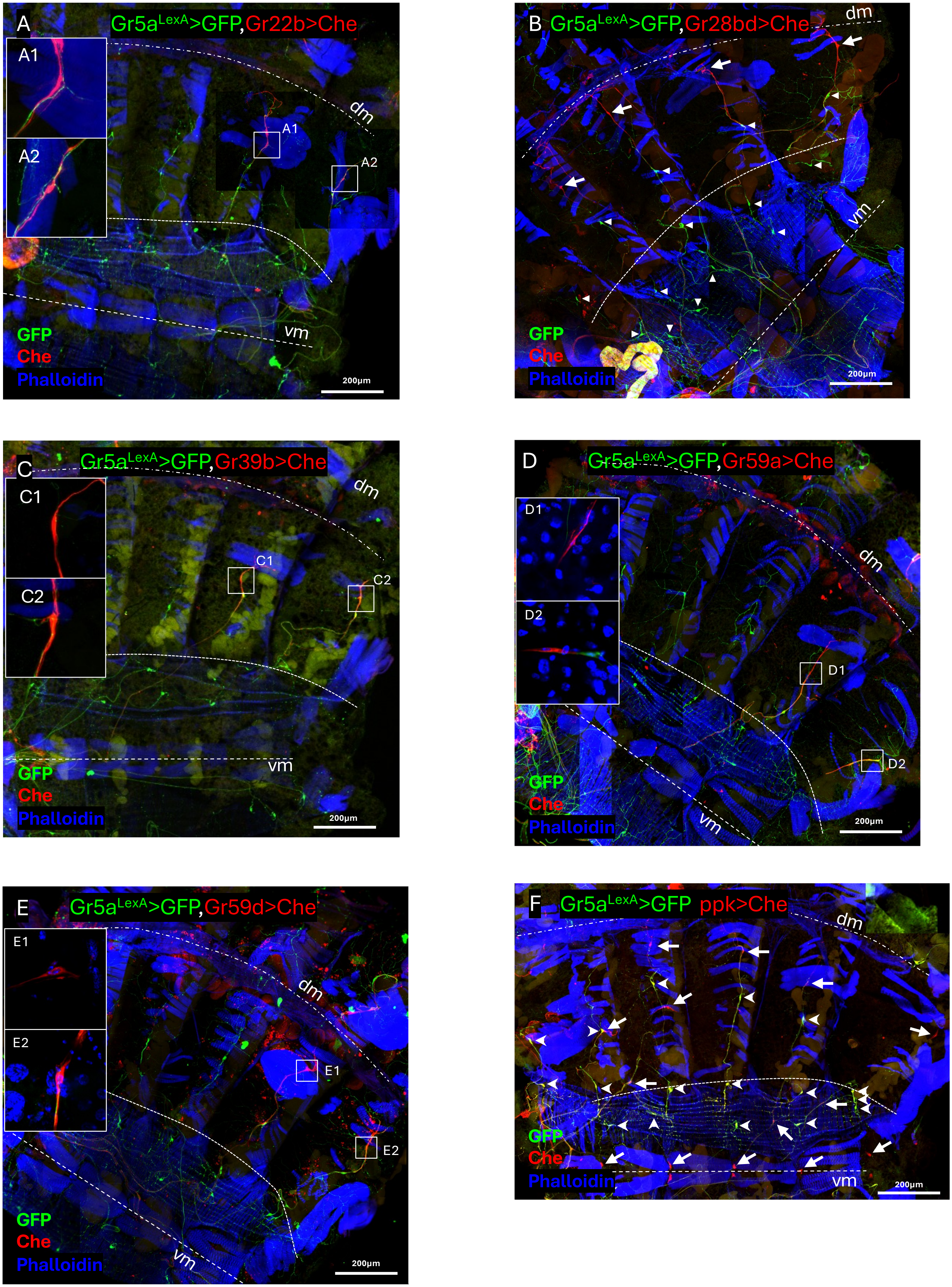
